# The contribution of uncharted RNA sequences to tumor identity in lung adenocarcinoma

**DOI:** 10.1101/2021.07.28.454105

**Authors:** Yunfeng Wang, Haoliang Xue, Marine Aglave, Antoine Lainé, Mélina Gallopin, Daniel Gautheret

**Affiliations:** Institute for Integrative Biology of the Cell (I2BC), Université Paris-Saclay, CNRS, CEA, 1 avenue de la Terrasse, 91190, Gif-sur-Yvette, France; Gustave Roussy, 114 rue Edouard Vaillant, 94800, Villejuif, France; Annoroad Gene Technology Co., Ltd,, 100176, Beijing, China

**Keywords:** k-mers, contigs, repeats, LUAD, mapping-free, replicability

## Abstract

**Background:** Transcriptome analysis of cancer tissues has been instrumental in defining tumor subtypes, diagnostic signatures and cancer regulatory networks. Cancer transcriptomes are still predominantly analyzed at the level of gene expression. Few studies have addressed transcript-level variations, and most of these only looked at splice variants. Previously we introduced a k-mer based, reference-free method, DE-kupl, that performs differential analysis of RNA-seq data at the k-mer level, which enables distinguishing RNAs differing by a single nucleotide. Here we evaluate the significance of differential events discovered by this method in two independent lung adenocarcinoma RNA-seq datasets (N=583 and N=154).

**Results:** Focusing on differential events in a tumor vs normal setting, we found events in endogenous repeats, alternative splicing and polyadenylation sites, long non-coding RNAs, retained introns and unmapped RNAs. Replicability was highly significant for most event classes (assessed by comparing to events shared between unrelated tumors). Overall about 160,000 differential k-mer contigs were shared between datasets, including a large set of sequences from hypervariable genes such as immunoglobulins, *SFTP* and mucin genes. Most interestingly, we identified a set of novel tumor-specific long non-coding RNAs in intergenic and intronic regions. We found that expressed endogenous transposons defined two major groups of patients (high/low repeat expression) with distinct clinical characteristic. A number of repeats, intronic RNAs and lincRNA achieved strong patient stratification in univariate or multivariate survival models. Finally, using antigen presentation prediction, we identified 55 contigs predicted to produce recurrent tumor-specific antigens.

**Conclusions:** K-mer based RNA-seq analysis enables description of cancer transcriptomes at nucleotide precision, independently of prior transcript annotation. Application to lung cancer data uncovered events stemming from a wide variety of transcriptional and postranscriptional mechanisms. Among those events, a significant subset was replicable between cohorts, thus constituting novel RNA hallmarks of cancer. The code is available at: https://github.com/Transipedia/dekupl-lung-cancer-inter-cohort.

## Background

Over a period of 20 years, cancer transcriptomics has transformed our understanding of tumor biology and led to improved tools for tumor typing and outcome prediction [1, 2]. While first generation transcriptome analysis was based on DNA microarrays with a focus on protein-coding genes, the current generation relies on RNA-seq data, which promises to deliver a more comprehensive view of gene expression. However, in spite of its potential for transcript discovery, cancer RNA-seq data is still utilized mostly to quantify the expression of annotated genes listed in a reference transcriptome. This ignores a wide array of mRNA isoforms, non-coding RNAs, endogenous retroelements and transcripts from exogenous viruses and bacteria [3]. The quantity of information left unexploited in non-canonical transcripts remains unknown. A number of studies have started to address this question using publicly available cancer RNA-seq data, focusing on specific transcript classes such as splice variants [4, 5], lncRNAs [6], snoRNAs [7], repeats [8], bacterial RNA [9], or viral RNA [10]. Other neglected sources of RNA diversity are the so-called blacklisted regions of the genome that are too variable or repeated to be properly analyzed by conventional approaches [11]. To our knowledge, no attempt has been made to extract and evaluate at once all this non-standard RNA information from tumor RNA-seq data. We think this approach could be particularly valuable in cancer since every individual tumor harbors a unique transcriptome that departs from that of normal tissues in multiple, unpredictable ways.

Previously we introduced a computational method, DE-kupl [12], that performs differential analysis of RNA-seq data at the k-mer level. As this method is reference-free and mapping-free, it identifies any novel RNA or RNA isoform present in the data at nucleotide resolution, including poorly mapped transcripts such as RNAs from repeats and chimeric RNAs. Here we set ourselves to evaluate all non-reference events discovered by DE-kupl in a comparison of normal vs. tumor samples using lung adenocarcinoma as a test case. To mitigate false positives events inherent to any gene expression profiling [13, 14], we focused on events that were replicated in two independent datasets. This required the development of a dedicated protocol to identify shared events in unmapped RNA sequences. Results revealed a collection of novel tumor-specific unannotated lincRNAs, intron retentions, and splicing events. Most strikingly, a collection of endogenous retroelements form a major class of tumor defining transcripts and constitute potent survival signatures. We also identified a subset of events with no expression in normal tissues which could be potential neoantigens sources. We would like to suggest DE-kupl as a promising, comprehensive approach to cancer transcript profiling.

## Methods

### Datasets

LUAD-TCGA: 582 lung RNA-seq samples from the LUAD-TCGA project were downloaded from the dbgap repository with permission, including 524 lung adenocarcinoma (LUAD) tissues and 58 adjacent normal tissues [15]. LUAD-SEO: The LUAD RNA-seq dataset of Seo et al. [16] was downloaded from the SRA database (accession: ERP001058). This dataset contains fastq files of 87 LUAD and 77 adjacent normal tissues. Only the 77 paired normal and tumor samples were analyzed. PRAD-TCGA: For control, 557 PRAD-TCGA prostate RNA-seq datasets were downloaded from dbgap with permission, including 505 prostate adenocarcinoma (PRAD) and 52 normal controls [17]. Bam format files from the TCGA datasets were converted to fastq format using Picard tools version 2.18.16 (http://broadinstitute.github.io/picard).

### DE-kupl pipeline

DE-kupl (version 5.3.0) was applied to the three datasets with the same parameters: in the filtering steps, k-mers with abundance fewer than 5 (min_recurrence_abundance) and present in no more than 10 samples (min_recurrence) were ruled out. In order to focus on non-canonical transcripts, we masked all k-mers pertaining to the main transcript of each Gencode gene as in [12]. Normalization factors for k-mer counts were computed by DE-kupl as medians of the ratios of sample counts by counts of a pseudo-reference obtained by taking the geometric mean of each k-mer across all samples. Herein we will use these counts as a proxy to represent the expression of the corresponding RNA fragment.

For differential expression analysis, the version of DESeq2 available at the time of the experiment was too slow for dealing with hundreds of samples and we found the faster “T-test” option to lack sensibility. Hence we used instead Limma [18], adapted to millions of k-mers using a chunk-based strategy (suppl. methods). This was found to perform 10 times faster than DESeq2. The performances of DESeq2, Limma and T-test for differential expression evaluation have been evaluated before [19]. Evaluations of k-mer counts were log-transformed and Limma was used to calculate log fold-changes and P-values. Retention thresholds for log2 fold changes and P-values were 1 and 0.05, respectively. All k-mers passing the filtering process above were merged into contigs and the contig table was saved as output. GC-contents in “up” and “down” contigs in the PRADtcga dataset were verified and did not present any bias (Additional file2: Table S1). High-quality contigs (”top contigs”) were contigs with counts>10 in at least 15% of the smaller class (Normal or Tumor).

Gene-level expression was measured using Kallisto v0.43.02 [20] and Gencode v31 transcripts, followed by summing TPM values of transcripts from the same gene. Gene-level differential expression analysis was performed using Limma and the same normalization procedure as above. Downstream analyses were conducted using R version 3.5.2. Heatmaps were drawn using the ComplexHeatmap package (version 2.4.3) [21].

### Shared event identification

Contigs from distinct DE-kupl analyses were decomposed into their constituent k-mer lists and a graph was constructed using the NetworkX Python package (version 2.3) [22], with k-mers as nodes and shared k-mers as edges. Contigs corresponding to the same local event are expected to form a fully connected subgraph or clique (Additional file 1: Fig. S1). We thus extracted all cliques to identify shared contigs. Hereafter we use the ∩ operator to represent contigs shared between two datasets.

### Contig annotation

A uniform annotation procedure was applied to contigs from each independent analysis (LUADtcga, LUADseo, PRADtcga) and to shared contigs (LUADtcga ∩ LUADseo and LUADtcga ∩ PRADtcga). Initially, differential contigs were mapped and annotated with DE-kupl annotation (https://github.com/Transipedia/dekupl). Briefly, DE-kupl annotation maps contigs to the human genome and reports intronic, exonic or intergenic status, CIGAR string, IDs of mapped or neighboring genes, differential usage status. A new repeat annotation field (“rep type”) was added based on Blast [23] alignments of contigs to the DFAM repeat database [24] (see Suppl. Methods). The results of DEkupl-annot were then loaded into R and submitted to further filtering and annotation. Firstly, a count filter was applied to retain only contigs with a count of 10 in at least 15% of the smaller class (Normal or Tumor). Contigs meeting this criterion were classified into event classes comprising SNV, intronic, splices, split, lincRNA, polyA, repeat and unmapped, as described in Additional file2: Table S3. Classes were non exclusive, meaning that a contig can belong to several classes. Since the TCGA datasets are unstranded, antisense events were not called. Differential usage (i.e. the relative change in expression of a local event relative to the expression of the host gene) was evaluated for each event mapped to an annotated gene. Intergenic contigs were further aligned with Blast against MiTranscriptome V2 [6] retrieved at http://mitranscriptome.org/ and converted to fasta using gffread (https://github.com/gpertea/gffread). Finally, we defined a new category called “neoRNAs”, which includes contigs that are expressed in tumor tissues but silent in normal tissues.

### Functional enrichment of intronic events

Candidate intronic events were identified based on the DE-kupl differential usage P-value (computed by comparing the expression or the contig with that of the host gene). Gene Ontology biological process enrichment of host genes was assessed using the clusterProfiler R package (version 3.16.0) [25].

### Sample clustering based on repeats

We used the K-means algorithm [26] to cluster LUAD patients into two main subgroups based on the expression of contigs matching AluSx, L1P1 orf2 and L1P3 orf2 repeats. Clusters were then analyzed for enrichment in clinical features, immune infiltration, tumor mutational burden and copy number variants. LUAD driver genes were retrieved from the COSMIC Cancer Gene Census (CGC) list [27]. Oncoplots were drawn using the maftools R package (version 2.4.10) [28]. The estimated tumor mutational burden (TMB) for each patient was computed using the total number of non-synonymous mutations from the Mutation Annotation Format (MAF) file, divided by the estimated size of the whole exome. Copy number variation (CNV) data was downloaded by the TCGAbiolinks R package (version 2.16.3) [29], which provides a mean copy number estimate of segments covering the whole genome (inferred from Affy SNP 6.0). The ratio of gain and loss for each patient was estimated by the fraction of segments indicating CNVs. Heatmap representations were produced with ComplexHeatmap [21].

### Correlation with immune infiltration

Immune infiltration analysis was performed on the LUADseo dataset. Relative proportions of infiltrating immune cells were determined using CIBERSORT [30]. Relationships between immune cell types and shared contigs (grouped by annotation category) were computed as the Spearman correlation between the contig expression and the relative proportion of the cell type in all samples. Any contig with an absolute Spearman correlation coefficient above 0.5 with at least one immune cell type was retained.

### Neoantigen prediction

For prediction of recurrent tumor-specific antigen, we selected contigs absent in all normal tissues but present in at least 15% of tumor tissues. We translated contig sequences using EMBOSS transeq over 6 frames [31]. Sequences with stop codons were ruled out and candidate peptides were submitted to netMHCpan 4.0 [32] to predict binding affinity to MHC-class-I molecules. Peptide–MHC Class I interactions with strong binding levels (by default 0.5%) were reported.

### Survival analysis based on event classes

Since the LUADseo dataset does not include survival information, we only performed the survival analysis on the LUADtcga dataset. Overall survival time and status was downloaded from the GDC portal (https://portal.gdc.cancer.gov/projects/TCGA-LUAD). We performed both univariate Cox regression and multivariate Cox regression on each event class to assess the prognosis value of the differential events. Survival analysis was performed using the survival (version 3.2.3) and survminer (version 0.4.7) R packages [33, 34]. Hazard ratios (HR) and P-values were calculated for each contig. Contigs with HR>1 and P-value<0.05 were considered as potential risk factors. For multivariate Cox regression, contigs were initially selected by cox-lasso regression using the glmnet R package (version 4.0.2) [35] applied independently to each contig class. The multivariate model was then constructed using selected contigs. Patients were divided into high and low-risk groups based on the median value of all risk scores for representation in Kaplan–Meier (KM) curves [36].

### Unsupervised clustering analysis

We applied Principal Component Analysis (PCA) and hierarchical clustering to each event class. PCA analysis was performed with the factoextra R package (version 1.0.7) [34]. Heatmap views were obtained using ComplexHeatmap [21].

### Sequence alignment views

We created “metabam” alignment files for tumor and normal tissues from each cohort. To this aim, we randomly sampled 1M reads from each fastq file of each subcohort using seqtk (https://github.com/lh3/seqtk) and aligned the aggregated reads to the genome (GRCh38) using STAR (version 2.7.0f) [37] with default parameters. BAM files were visualized using Integrative Genomics Viewer (IGV 2.6.2) [38].

## Results

### Gene-level *vs*. contig-level differential events

We performed tumor *vs*. normal differential expression (DE) analysis on two independent Lung adenocarcinoma RNA-seq datasets from TCGA (LUADtcga) and Seo et al. (LUADseo) and on a prostate adenocarcinoma dataset from TCGA (PRADtcga) as a control. Each dataset was submitted to a conventional, genelevel, differential expression analysis and a k-mer level differential expression analysis where all k-mers from annotated genes were first removed and the resulting differential k-mers were assembled into contigs (Fig 1A). For simplification, we shall hereafter use term “expression” when referring to either gene expression or contig k-mer counts. While the number of DE genes in the three comparisons ranged from 6,000 to 9,000, the number of DE k-mers was about a thousand times larger (2 to 12 millions). Assembly of k-mers into contigs reduced this number to about 400,000 DE contigs in each analysis (Fig 1B).

**Figure 1.**
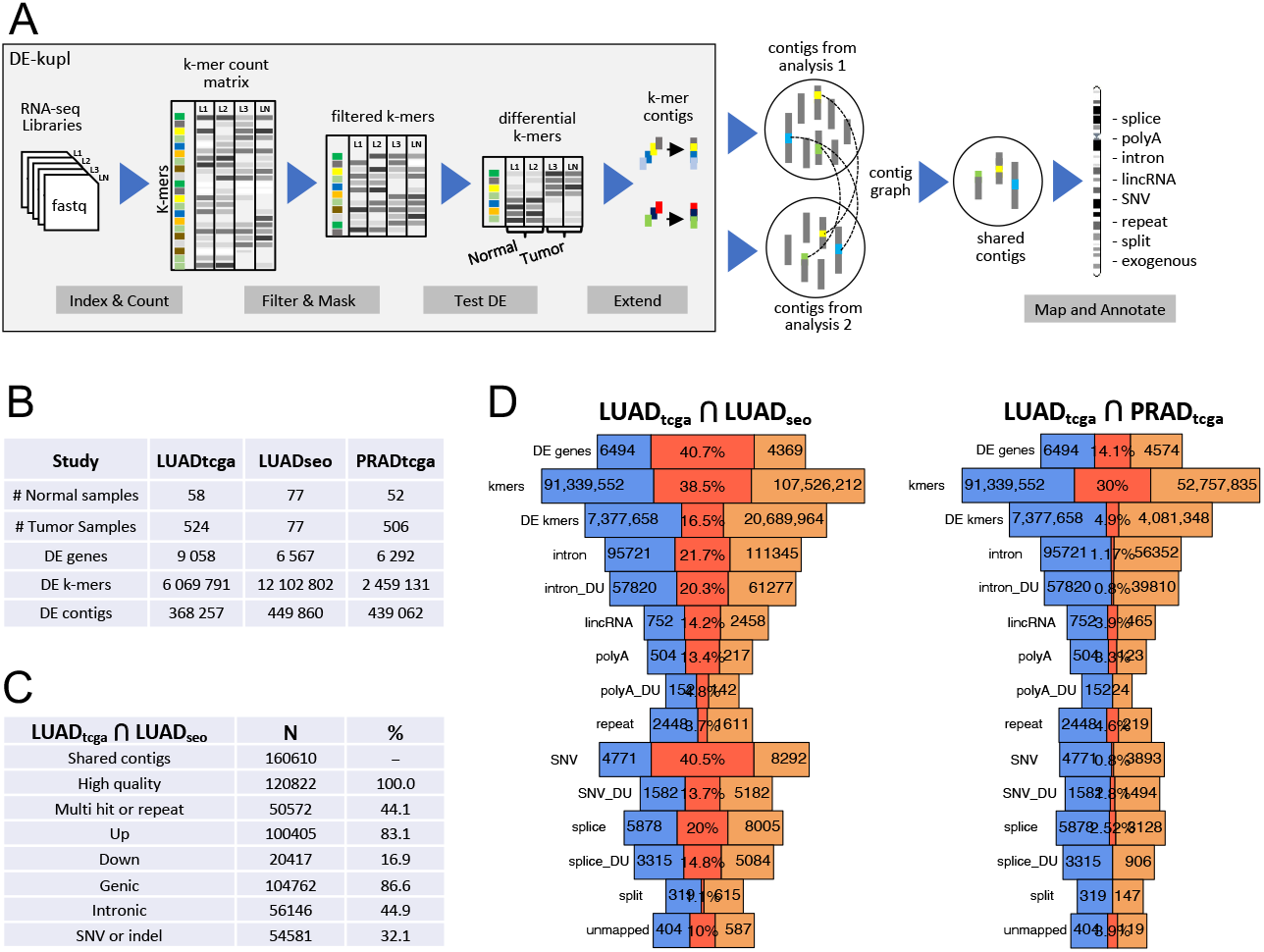
Overall analysis procedure and properties of identified contigs. (A) Computational pipeline for inferring differential contigs in each tumor/normal cohort, extraction of shared contigs and annotation. (B) Sizes of RNA-seq cohorts analyzed and numbers of differential events observed. (C) Summary statistics of differential contigs identified as shared between the LUADtcga and LUADseo analyzes. (D) Number of differential genes, k-mers and contigs in each independent analysis and shared between analyzes. On each row, lateral areas represent differential genes/k-mers/contigs found in each independent analysis and the central area represents shared differential genes/k-mers/contigs. Contigs are classified into different annotation groups.

We next compared the DE genes and contigs discovered in independent datasets to identify shared DE events. While this process is trivial for genes, it is not for contigs, since contigs found in each dataset have no standard identifier that could be used to relate them. We thus implemented a graph analysis procedure that identified shared contigs based on their common k-mers (Fig 1A, Additional file 1: Fig. S1). A final annotation step assigned contigs to non exclusive categories based on their mapping characteristics or expression (repeats, lincRNAs, splice variant, polyadenylation variants, split RNAs, tumor-specific RNAs) as described in Additional file2: Table S3 and Methods. The numbers of shared elements slightly differ between LUADtcga and LUADseo because a minority of elements are in a 2-to-1 or 1-to-2 relationship in the contig graph. If not otherwise specified, numbers of elements are given for the LUADtcga cohort.

Overall 160,610 differential contigs were shared between the two LUAD analyses (Fig 1C). Over these, 120,822 contigs were considered of sufficient quality based on counts and occurrence in a minimal number of samples (see Methods). 83% of shared contigs were overexpressed in tumors vs. only 17% underexpressed (Fig 1C).

### Event replicability

The replicability of differential events was generally lower for k-mer or contigs than for genes. Fig 1D shows the number of differential expression genes and contigs shared by the two independent LUAD analyzes, with contigs binned by annotation class. About 41% of differential expression genes (3032 genes) were shared by the two LUAD analyses, compared to an average of 14% for differential expression contigs (repeats: 3.7%, unmapped RNAs: 10%, alternative polyAs: 13%, lincRNAs: 14%, alternative splices: 20%, retained introns: 20%). Although the ratio of shared events was relatively low for k-mer analysis, it was considerably higher than when comparing two unrelated pathologies (LUADtcga ∩ PRADtcga, Fig 1D), and this applied to all event classes except repeats. This indicates that, although k-mer based differential expression events are noisy, a significant subset is replicable in independent studies. Furthermore, we observed a strong correlation between the fold-change value of differential expression contigs and the likelihood to be shared between cohorts (Additional file 1: Fig. S2), demonstrating the non-randomness of high scoring, non-reference events.

### DE contig localization, hypervariable genes

The majority of shared contigs are genic (83%), 45% are intronic and 32% carry SNVs or indels (Fig 2A). These characteristics are induced by the initial filter that removed all k-mers matching reference transcripts, retaining any intronic or SNV-carrying k-mer. Therefore a large number of SNV and intronic contigs are just “passenger” events of DE genes. We confirmed this by analyzing the correlation between numbers of DE contigs and host gene expression. We found a significant correlation (Pearson CC=0.45), but this correlation was reduced (Pearson CC=0.28) in shared DE contigs, indicating shared contigs contain fewer passenger events (Additional file 3).

**Figure 2.**
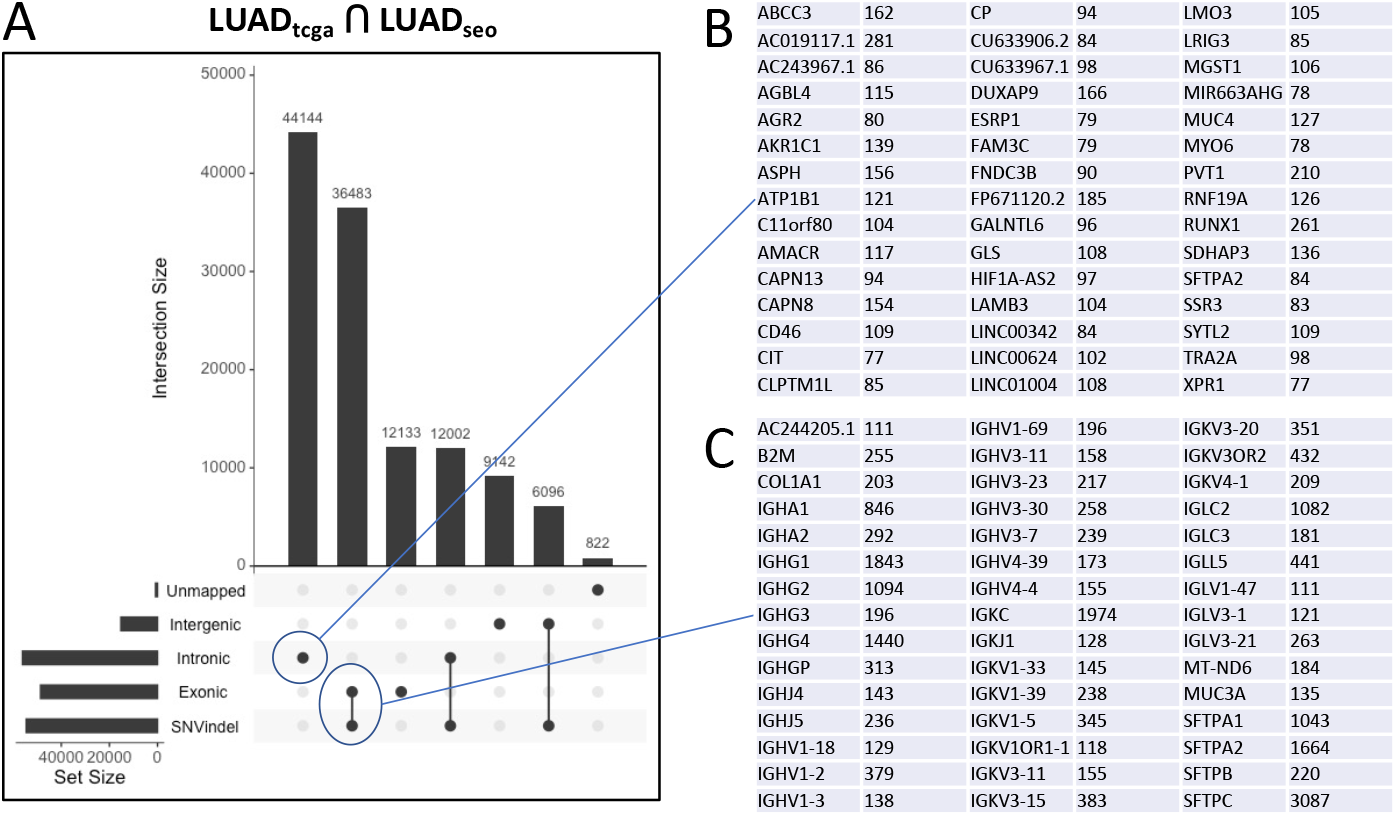
General properties of shared differential expression contigs in LUAD. (A) UpsetR plot of major contig categories based on mapping location and presence of SNV or indels. (B) 45 top genes by number of mapped contigs in the circled intronic category. (C) 45 top genes by number of mapped contigs in the circled exonic+SNVindel category. Numbers of contigs mapped to each gene are indicated.

More than 400 genes were matched by 35 or more contigs. We classified these genes into two categories: for 296 genes, most contigs matched introns and were up-regulated in tumors (Fig 2A, B, Additional file 2: Table S5). These mostly correspond to the aforementioned passenger events. The second category is composed of 107 genes we refer to as “hypervariable” as they tend to yield a large number of contigs carrying SNVs, indels and larger rearrangements (Fig 2A, C, Additional file 2: Table S5). The largest sets of hypervariable genes are *IGK*, *IGL* and *IGH* immunoglobulin genes. This is not surprising given immunoglobulins (i) are highly variable due to V(D)J segment recombination and (ii) are expressed by plasma B-cells which are abundant in the tumor immune infiltrate [39], hence these genes are seen as up-regulated in tumors. Interestingly, those IG sequence variants are found expressed in different patients and across the two cohorts, suggesting our approach can be used to profile immunoglobulin repertoires, as performed recently with other RNA-seq datasets [40]. To evaluate the accuracy of DE-kupl contigs assembled from IG genes, we selected all contigs mapped to one arbitrary IG gene (IGHV: 100 contigs) and aligned them to IGHV contigs from the IMGT database [41]. Ninety out of 100 contigs had significant matches in the corresponding IMGT category extending over 90% of the contig length (Additional file2: Table S6).

Other hypervariable loci were found in surfactant protein (*SFTP*) and Mucin genes which are known to harbor a high level of polymorphism [42, 43]. We observed polymorphism not only in the form of SNPs, but also in the form of splicing variations. Five *SFTP* genes alone combine over 9000 SNVs and 800 splice sites contigs, while 12 Mucin genes harbour 1324 contigs including 42 splice variants (Additional file 1: Fig. S3A-B, Additional file 2: Table S5). While *SFTP* contigs were all underexpressed in tumors, Mucin contigs were mostly overexpressed (Additional file 2: Table S5). Mucins are immunogenic [43] and are important biomarkers for prognosis [44] and drug resistance [45]. The existence of recurrent mucin variants overexpressed in tumors may be relevant for these therapeutic and biomarker developments. We also observed hypervariability in *CEACAM*5 and *KR*19, two other prognostic biomarkers and/or immunotherapy targets [46, 47] (Additional file 1: Fig. S3C, Additional file 2: Table S5).

### Intron retention and other intronic events

We found intronic contigs with differential usage (DU) in 313 host genes, 290 (93%) of which were up-regulated in tumors (Additional file 2: Table S4). 70% of the host genes were also up-regulated, thus the apparent overexpression of these intronic sequences may have been confounded by overexpression of host genes. However, 30% of host genes were not overexpressed, and in 103 cases, intron and host gene expressions varied in opposite directions (93 introns up and 10 introns down). Our annotation pipeline did not differentiate intron retentions (as shown for example in Additional file 1: Fig. S4A) from transcription units occurring within introns (example in Additional file 1: Fig. S4B). We observed intron retention events in lung cancer drivers *EGFR* and *MET* (Additional file 1: Fig. S4C and Additional file 1: Fig. S4D). In *EGFR*, the retained intron was located between exons 18 and 19, just upstream of the principal oncogenic *EGFR* mutations located in exons 19-21. Intron retention before exon 19 would likely produce a truncated form of *EGFR* compatible with oncogenic activation.

Additional file 1: Fig. S5A shows the 20 intronic events with the most significant differential usage P-values. All show opposite directions of intron and gene expression. Gene Ontology enrichment analysis indicates host genes are enriched for inflammation and immune response pathways involving neutrophil and T cells (additional file 1: Fig. S5B), suggesting these events may come from regulations in the tumor microenvironment rather than in the tumor itself.

### Novel lincRNAs

Contigs that do not map any Gencode annotated gene are of particular interest as they potentially represent novel lincRNA biomarkers of lung tumors. Overall we identified shared DE contigs in 885 intergenic regions, which we labelled as lincRNAs. As genic regions already included annotated lncRNAs and pseudogenes from Gencode, the actual number of DE contigs in lncRNAs and pseudogenes was much higher (N=2892) but we focus here on unannotated regions. lincRNA contigs were mostly overexpressed in tumors (83% of contigs) and often contained a known repeat element (73% of contigs). Their average length was 137 nt, however actual transcription units were generally longer as most units were composed of multiple contigs, as shown in examples in Additional file 1: Fig. S6. Most intergenic contigs (793 out of 823) were already annotated in the independent Mitranscriptome lncRNA database [6], which was expected since this database was also produced from TCGA RNA-seq data. Less than one third of the flanking genes of intergenic contigs were differentially expressed, indicating that novel lincRNA expression was most often independent from that of flanking genes.

### Expressed repeats delineate patient subgroups with distinct clinical properties

The dominant model for endogenous retroelements (EREs) expression is that EREs are mainly expressed in germline and embryonic stem cells while they are repressed in differentiated somatic cells. However recent studies have shown expression of EREs in somatic cells is more common and heterogeneous than expected[48]. Repeat-containing reads are difficult to analyze by RNA-seq standard pipelines due to ambiguity in the alignment process. We thus questioned whether our alignment-free procedure could help reveal these events. From the initial set of 50572 contigs annotated as repeats (Fig 1C), we selected a high quality subset of 10341 contigs over 60 bp in size and with expression above a set threshold (see Methods). Of these, 87.7% were overexpressed in tumors (Additional file 2: Table S4).

Fig 3A shows the distribution of contigs per repeat family. Most repeats correspond to Line 1 and Alu family sequences. The most frequent repeat overall is L1P1, a Line 1 of the L1Hs family which is the only retrotransposition-competent EREs in the human genome [49]. L1P1/L1Hs elements, as well as human endogenous retrovirus (HERV), were almost exclusively over-expressed in tumors, suggesting tumor-specific activation of these elements. In contrast, Alu elements, which are often expressed as part of protein coding genes, were either over- or under-expressed in tumors. Fig 3A shows the top 20 repeat types that contribute more contigs. Fig 3B-C shows the expression heatmap of the 60 repeats contributing more contigs. For each type of repeats, we selected the contig with the highest absolute fold-change.

**Figure 3.**
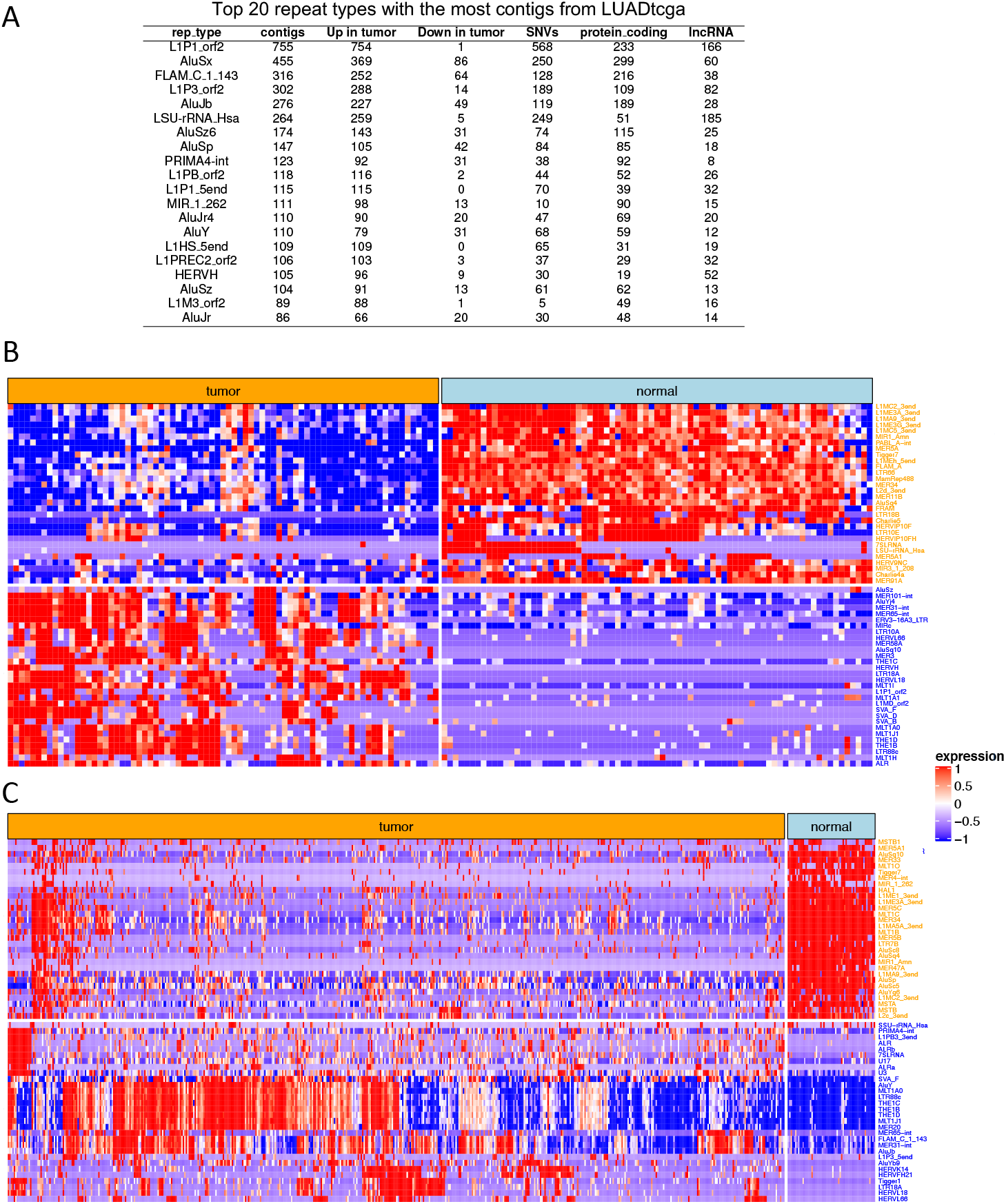
Tumor identity based on top repeats (A) Top 20 repeat types with the most contigs in LUADtcga dataset. (B/C) Expression heatmap of top repeat-containing contigs (ranked by fold change) in the LUADseo (A) and LUADtcga (B) datasets. Contig expression level in heatmaps is represented from blue (lowest) to red (highest).

Repeat contigs also included a group annotated as “simple repeats”, containing microsatellites and other low complexity elements. Contrarily to EREs, these do not have the capacity to be expressed independently. Indeed, in over 70% of cases, these contigs were uniquely mapped to genic sequences. In addition to annotated repeats and simple repeats, DE-kupl identified 4762 contigs (4497 up, 265 down) with multiple genome hits but no match in the DFAM repeat database (Additional file 2: Table S4). Many of these repeats were from Mucins, immunoglobulins and multicopy gene families such as *NBPF* and *TBC*1. These repeats are shared between two cohorts and thus represent robust events of (mostly) overexpressed RNA fragments in tumors that would hardly be noticed in regular RNA-seq analysis due to their low mappability.

To investigate repeat-based patient subgroups, we performed clustering of tumors based on the most frequent repeat elements in Fig 3A: AluSx, L1P1 orf2, and L1P3 orf2 (as FLAM repeats are a family of Alu-like monomers that give birth to the left arms of the Alu elements, we did not account for FLAM C 1 143). K-means clustering with *k* varying from 2 to 4 groups consistently found two major subgroups: subgroup 1 (“repeat-low”) displayed generally low expression of Alu and L1 repeats compared to subgroup 2 (“repeat-high”) (Fig 4A).

**Figure 4.**
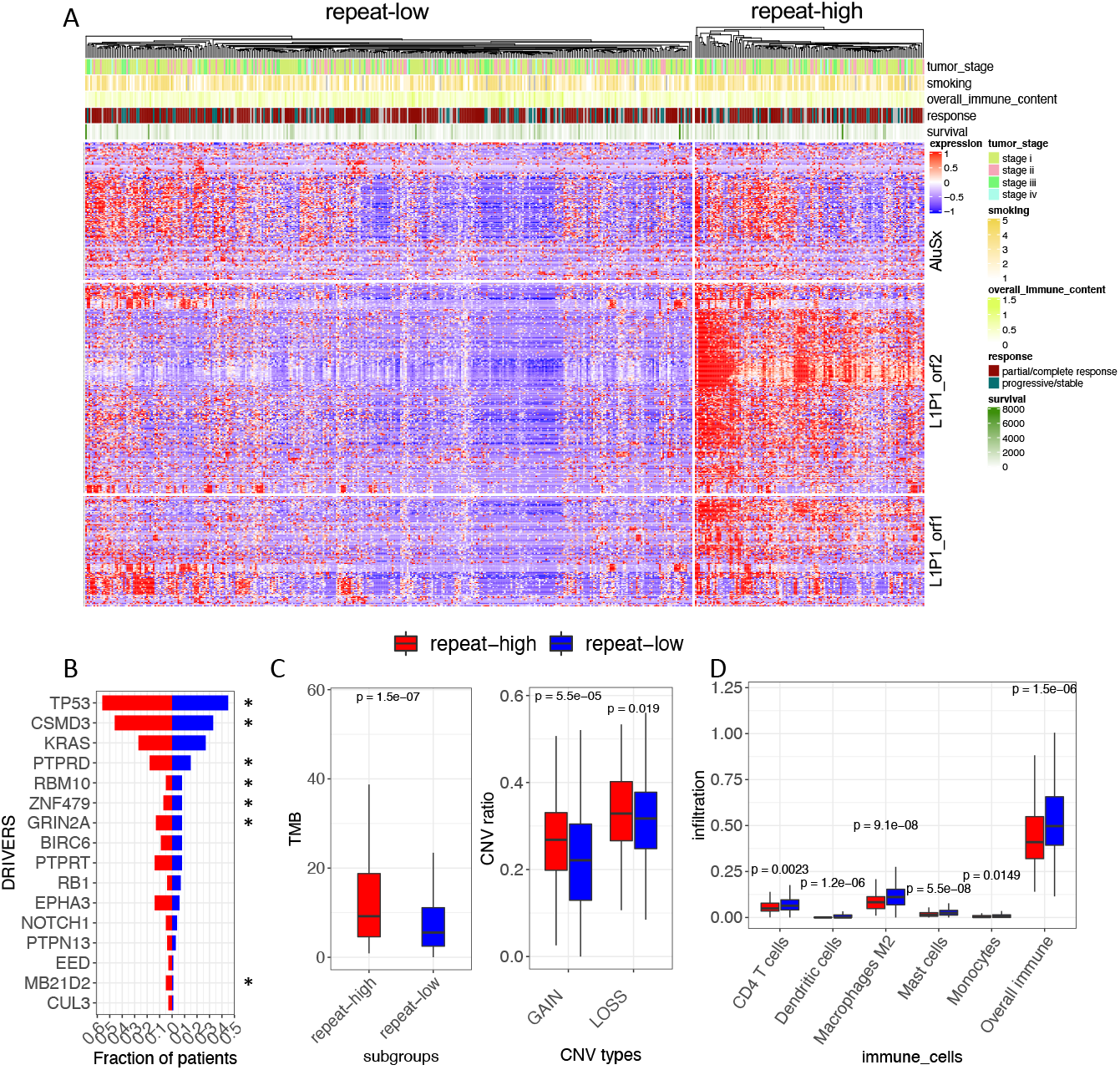
Characterization of patient subgroups based on repeat-containing contigs. (A) Clustering of LUADtcga patients into two subgroups based on Alu and L1P1 repeat expression. Subgroups were defined by K-means. (B) Fraction of patients with driver mutations for 16 COSMIC LUAD drivers. Drivers with Fisher P value < 0.05 were marked with star. (C) Mutational burden and CNV frequency distribution between two subgroups. (D) Variation of immune features between subgroups. The red and blue represent the repeat-high and repeat-low subgroups, respectively. P-values are computed by Wilcoxon test.

We then related the two repeat subgroups with somatic alterations observed in TCGA patients. Patients in the repeat-high group were more frequently mutated in LUAD drivers *CSMD*3, *TP*53, *PTPRD*, *PTPRT*, *GRIN*2*A*, *EPHA*3, and *MB*21*D*2 (Fig 4B, Fisher P<0.05). Patients in the repeat-high group had a significantly higher TMB (Wilcoxon P=1.5e-07) and a higher ratio of CNVs than other patients (Wilcoxon P=5.5e-05 for gain; P=0.019 for loss) (Fig 4C).

We observed no difference between subgroups in terms of age, gender, tumor stage, overall survival (OS), and vital status, but found more smokers in the repeat-high group (Wilcoxon P=0.02). We then assessed the immune cell contents of samples estimated by gene expression deconvolution. The repeat-high subgroup had lower proportions of dendritic cells, M2 macrophages, mast cells, monocytes and CD4+ T cells and overall immune content than the repeat-low subgroup (Fig 4D). In summary, “repeat-high” tumors associate with higher genome instability, more frequent smoking and lower immune infiltration.

### Immune cell-associated contigs

We sought which contigs best correlated with tumor immune cell contents estimated by gene expression deconvolution. Sixty five contigs were found correlated with at least one type of immune cell (Additional file 1: Fig. S7). Most of these were uniquely mapped to genic introns or exons and underexpressed in tumors. Positive correlations were mostly observed with M2/M0 macrophages or resting CD4+ T cells, *i.e*. with a generally repressive or quiescent immune environment. However, a few contigs were associated to immune active M1 macrophages, including two contigs matching *GBP*5 (a marker of activated macrophages) and *CXCR*2*P*1 (a pseudogene expressed in an intron of *RUFY*4, a gene expressed in dendritic cells). Overall, immune cell-associated contigs mapped leukocyte-specific or immunity-related genes, suggesting most contigs originated from the immune cell themselves (Additional file 2: Table S11).

Perhaps the most intriguing set of immune cell-associated contigs was that correlated to naive CD4+ T-cells. These cells are not especially enriched in tumor or normal samples, yet they correlate with six DE contigs. One contig was strongly repressed in tumors and corresponded to *Klebsiella pneumoniae* large subunit rRNA. Indeed, *Klebsiella* is a common lung bacterium against which cross-reactive T-cells are present in the naive CD4+ T-cell repertoire [50]. Our results thus suggest the joint occurrence of *Klebsiella* and matching CD4+ T-cell in normal lungs, and their disappearance in tumors. Of note, this *Klebsiella* contig also correlates positively with multiple contigs in the *SFTP* gene (Additional file 2: Table S12), in line with *SFTP* roles in defense against respiratory pathogens [51].

The other five contigs associated with naive CD4+ T-cells were all overexpressed in tumors. These included two intergenic repeats related to HERV (human endogenous retrovirus): HERV-E and MER9. The HERV-E contig was expressed from the *env* gene of a near full-length retroelement. One may hypothesize that expression and antigen production by the *env* gene trigger recruitment of CD4+ T-cells, as observed already in breast cancer [52]. Alternatively, reactivation of HERV elements could be an intrinsic feature of the CD4+ T-cells [53]. This analysis illustrates how non-reference RNA quantification can illuminate the interplay between cell types and specific RNA elements including exogenous elements in a bulk tissue.

### Novel sources of shared neoantigens enriched in lincRNAs

Tumors express a large diversity of transcripts that are not usually expressed in normal tissues. When translated, these transcripts can produce peptides recognized as non-self by the epitope presentation machinery, triggering antitumor immune response [54]. These tumor-specific antigens or neoantigens are the object of active investigation for immunotherapy and tumor vaccine development. Protocols for neoantigen discovery usually start from a list of nonsynonymous somatic mutations identified from WES or WGS libraries and whose expression is confirmed by RNA-seq. Candidate mutated peptides are then submitted to an epitope presentation prediction pipeline [55]. This protocol predicts potential neoantigens from annotated and mappable regions. However, neoantigens can be produced from any transcript, including repeats and supposedly non-coding lncRNAs [56, 57]. Therefore we thought our reference-free approach could be a good source for such elements.

We considered contigs with no expression in normal tissues as potential neoantigen sources. To focus on shared neoantigens, we further requested contigs to be expressed in at least 15% of tumor samples. This selected 2375 contigs in the LUADtcga dataset (Fig 5.A). About 20% of these contigs (N=472) where also silent in normal tissues of the LUADseo cohort (Fig 5.B). We evaluated the potential of these “strictly tumoral” contigs for neoantigen presentation. Fifty five strictly tumoral contigs produced peptides predicted to be strong MHC-class-I binders by netMHC-pan (Additional file 2: Table S10). Although potential neoantigen-producing contigs were found in several categories and locations, intergenic location was the most significantly enriched category (Additional file 1: Fig. S8). Overall, contigs from intergenic regions, non-coding RNAs and pseudogenes contributed 58% of predicted neoantigens (Additional file 2: Table S10), consistent with previous reports of abundant neoantigen production from non-coding regions in other cancers [57].

**Figure 5.**
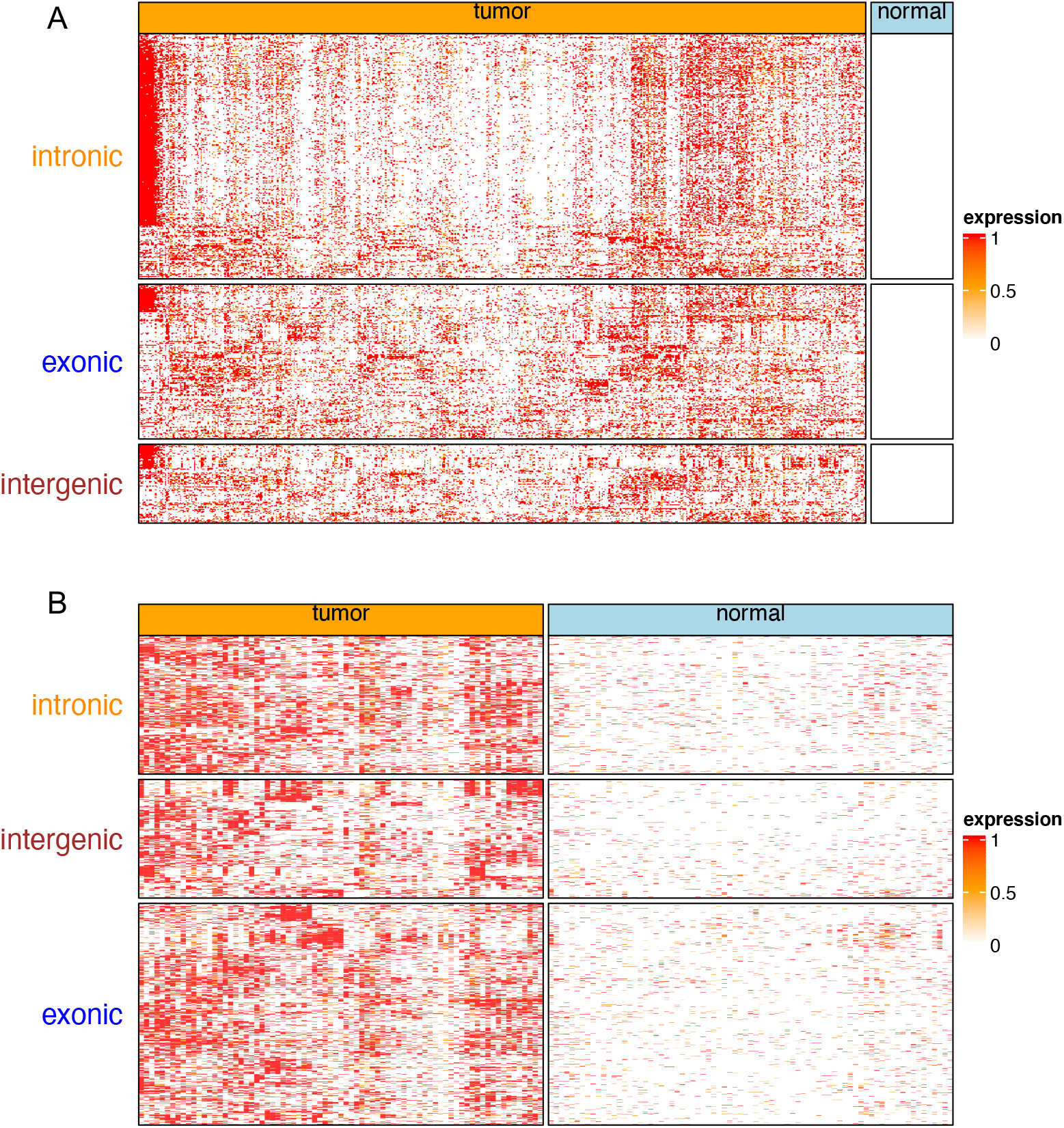
Expression heatmap of potential neoantigen sources in the two LUAD datasets. Tumor-specific contigs were first selected in LUADtcga (A) and validated in the LUADseo dataset (B).

### Repeats, intronic RNAs and lincRNA as survival predictors

To identify RNA elements associated with outcome, we retrieved overall survival (OS) data for the TCGA cohort and performed univariate Cox regression with the different classes of contigs. Thirty nine contigs were significantly related to OS after multiple testing correction (Additional file 2: Table S7). Outcome-related contigs are mostly enriched in repeats (Additional file 2: Table S8), especially HERV elements (4 out of the 10 top repeats) and Alu/L1 family elements (AluSx and L1P3 orf2). While HERV elements expression was always negatively related to OS, the trend for other repeats was variable, with different Line1 and Alu elements having either positive or negative relation to OS (Additional file 2: Table S7). Another interesting OS-related element was a novel splice variant in ELF1, a transcription factor of the ETS family involved in multiple cancers (Additional file 2: Table S7)[58].

We then performed multivariate Cox regression using sets of contigs selected by lasso regression within each contig category and using differentially expressed genes (Additional file 2: Table S9). Models based on annotated and simple repeats had the best prognostic power (log-rank P=2e-16, 2e-13, respectively, Fig 6). The “annotated repeat” model was based on 12 contigs, including six L1 and three HERV elements, reinforcing the relevance of these repeats for prognosis. The “simple repeat” model included 12 contigs with microsatellite-like repeats, of which 11 were uniquely mapped to the genome (Additional file2: Table S9). Other strong outcome predictors were obtained using lincRNA, intronic and unmapped contigs, all of which achieved a better patient stratification than a model based on DE genes (Fig 6).

**Figure 6.**
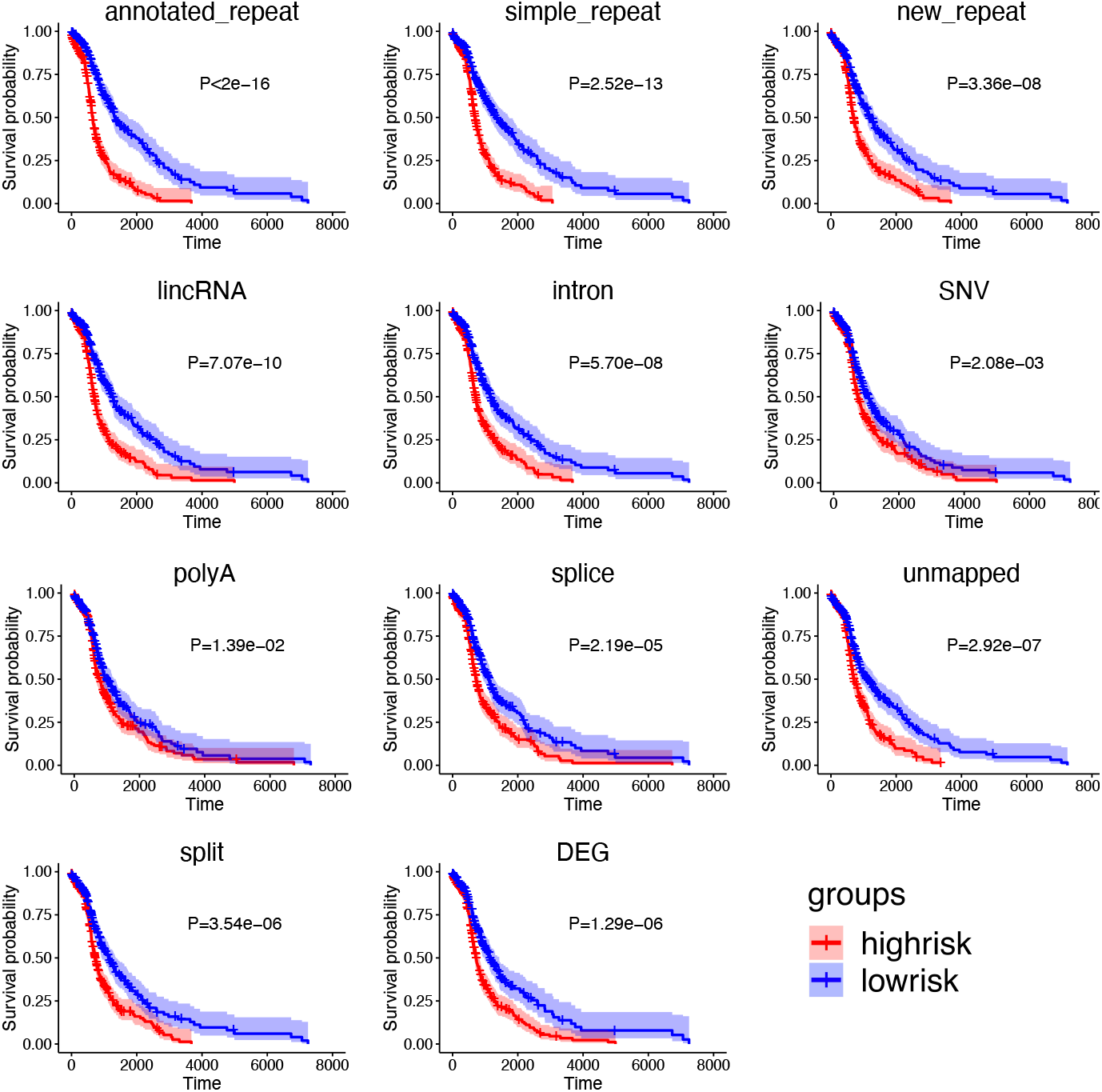
Kaplan-Meyer curves of multivariate survival models per class of event. Patients in high and low-risk groups are shown in red and blue, respectively. Repeat events were separated into annotated, new and simple repeats. The other categories with more lasso-selected contigs were also included (Additional file 3: Table S8).

### Unsupervised sample clustering based on non-reference RNAs

To investigate the capacity of non-reference RNAs to distinguish tumor and normal tissues in an unsupervised fashion, we performed PCA clustering of samples using contigs from each class (Fig 7). Tumor and normal tissues can be distinguished based on SNV, splice, intron, and lincRNA event classes as clearly as based on differentially expressed genes (“DEG” in Fig 7). This capacity is consistently observed in both cohorts. However, while many repeats are important with respect to tumor subclasses and survival, repeats altogether do not permit a clear separation of tumor and normal tissues in unsupervised clustering. Classes “polyA”, “split” and “unmapped” did not achieve clear separation either, which was more expected as these sets were much smaller in size.

**Figure 7.**
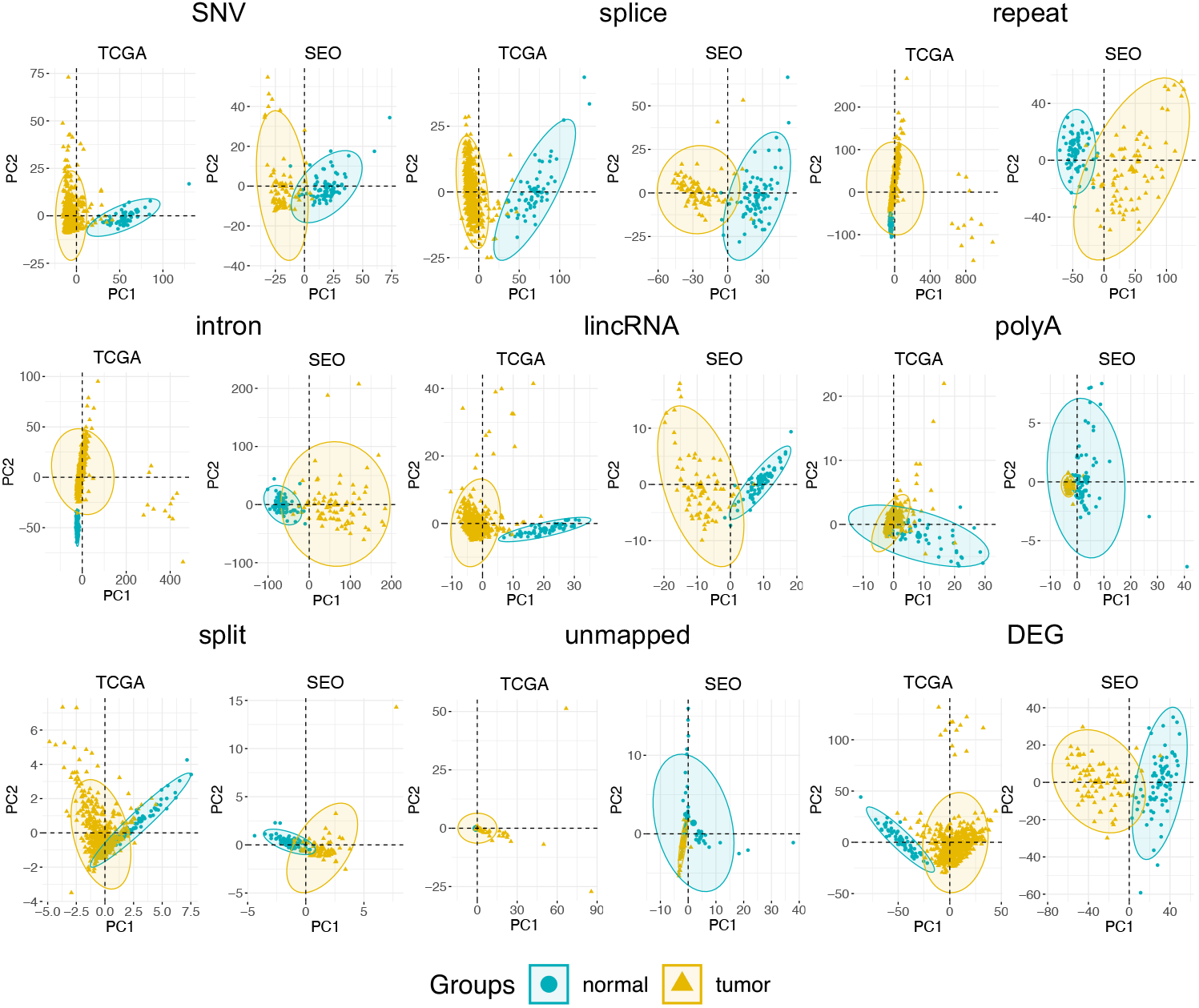
Principal component analysis of samples based on DE contigs and genes. Each panel represents PCA performed with one class of contigs and with differentially expressed genes (DEG), for the LUADtcga (TCGA) and LUADseo (SEO) datasets. Normal and tumor samples are marked using blue and yellow, respectively. Confidence ellipses are drawn with package factoextra for each group.

## Discussion

Using reference-free analysis of LUAD RNA-seq data, we identified a large set of differential RNA elements that were present in two independent LUAD cohorts. We classified these elements based on their genomic location, mapping characteristics and repeat contents. We did not analyze in detail all contig classes but focused instead on contigs mapping to hypervariable genes, repeats, lincRNAs and intronic elements. Besides these, a number of splice variants, chimeras, exogenous (non-human) sequences were found differentially expressed and could be pursued further.

A defining class of differential events involved endogenous repeats. The expression of L1 and Alu repeats defined two major tumor subgroups. The subgroup with higher L1/Alu expression was associated with more frequent mutations in *P*53, a higher mutational and copy number burden and a reduced immune cell infiltrate. This is consistent with previous observations that retrotransposition events can be controlled by *P*53 [59], correlate with a repressed immune environment [59, 60] and can lead to genome instability [61]. Expressed repeats also had significant prognostic power. Multivariate signatures composed of HERV and L1 elements, or simple repeats, stratified patients into distinct survival groups. Of note, HERV expression has been sporadically involved in various cancer types [62] and has recently been associated with poor prognosis in colorectal cancer [63].

A limitation of k-mer approaches for TE analysis is that transcripts are not fully assembled and thus the nature of repeats, whether expressed as functional retroelements or as part of mRNA or lncRNAs cannot be systematically established. Nonetheless, the majority of DE contigs are long enough to enable unambiguous mapping on the human genome, hence their origin could be further explored, including when coming from novel insertion events.

An attractive aspect of reference-free RNA-seq analysis is the capacity to identify novel forms of known cancer drivers or biomarkers. Indeed, we identified novel intron retention events in *EGFR* and *MET* and multiple new variants of *CEACAM*5 and *KR*19. Perhaps even more interesting is the ability to detect potential neoantigen sources in variant transcripts. Tumor-specific neoantigens have previously been identified from repeats and non-coding regions using mapping-based strategies [54, 57]. However, our approach casts a wider net as it collects all events independently of their origin, including when arising from unmappable or profoundly rearranged regions. Indeed we identified about 500 strictly tumoral contigs shared by patients from the two independent cohorts, 55 of which were predicted to produce MHC-class-I neoantigens. These shared neoantigen candidates are of particular interest since their targeting by antitumor therapy would potentially benefit groups of multiple patients.

The wealth of information uncovered in the present study is a strong incentive to explore other applications of reference-free transcriptomics. One such application is the identification of patient-specific abnormal transcripts under a 1 *vs n* experimental design, which is addressed by the Mintie software [64]. Reference-free strategies can also be used for building predictive models. We [65] and others [66, 67] are exploring this kind of approach to classify cancer RNA-seq samples with promising results. Finally, reference-free differential analysis of the type used in this study could be of particular interest in meta-transcriptomics projects where RNAs are sequenced from an environment containing unknown bacterial, archaeal or eukaryotic species. Our protocol guarantees that any RNA that is specific to a sample subset will be captured independently of its origin. We hope the present analysis will encourage others to explore other data sources in a reference-free manner.

## List of abbreviations

SNV: Single-Nucleotide Variants
CNV: Copy Number Variant
SV: Structural Variant
AS: Alternative Splicing
TCGA: The Cancer Genome Atlas
LUAD: Lung Adenocarcinoma
PRAD: Prostate Adenocarcinoma
EREs: endogenous retroelements

## Declarations

### Ethics approval and consent to participate

Not applicable

### Consent to publish

Not applicable.

### Availability of data and materials

Not applicable.

### Authors’ contributions

YW and DG designed the workflow and analyzed the results, YW downloaded and processed the datasets, YW and DG wrote the manuscript, MA and MG assisted in statistical analysis, HX assisted in coding scripts. AL annotated the repeat types.

## Acknowledgements

The results shown in this work are in whole or part based upon data generated by the TCGA Research Network: https://www.cancer.gov/tcga.

## Additional files

Additional file 1 — Figure S1

The graph-based protocol detecting shared contigs between TCGA and SEO datasets. (A) Contigs from each dataset. The bars marked with the same color represent the same k-mers. (B) Cliques construction based on the common k-mers. (C) Shared contigs identification based on the cliques.

Additional file 1 — Figure S2

Enrichment analysis of shared DEGs and contigs between TCGA and SEO datasets. The x axis represents the ranked DEGs or contigs based on log2FC in ascending order. The red vertical dotted line represents the position of log2FC cutoff.

Additional file 1 — Figure S3

Hypervariable genes in our analysis. Upset graph shows the overlap between different categories, including intronic, exonic, spliced and SNV or indel.

Additional file 1 — Figure S4

IGV views of intronic events. Each frame shows a metabam file composed of randomly sampled reads corresponding to the subcohort indicated on the left panel. The lower panel shows DE contigs and Gencode annotation. A: multiple intron retention in CEACAM5; B: lncRNA element expressed in an intron of MBD5; C: intron retention in EGFR; D: intron retention in MET.

Additional file 1 — Figure S5

Intronic event analysis. (A) Log2FC values of the top 20 intronic events (DU). Red and blue colors represent the expression fold change of intronic contigs and host genes, respectively. (B) Gene Ontology functional enrichment. Color represents the P-values and size represents the ratio of genes.

Additional file 1 — Figure S6

IGV views of lincRNA elements overexpressed in tumors. Each frame shows a metabam file composed of randomly sampled reads corresponding to the subcohort indicated on the left panel. The lower panel shows DE contigs and Gencode annotation.

Additional file 1 — Figure S7

Heatmap of Spearman correlation coefficient (CC) of contig counts and abundance of immune cell types evaluated by CIBERSORT. All contigs with a CC>0.5 with at least one immune cell type are shown. Immune cells not correlated with at least one contig are not shown. Row names show gene symbols and repeat types of contigs, whenever applicable. Row name colors indicate different contig categories. The log2FC sidebar shows expression fold change of contigs between normal and tumor samples.

Additional file 1 — Figure S8

Fractions of event types in strictly tumoral contigs predicted to produce neoantigens (“neo”, N=472) and total shared DE contigs (“all”, N=2375). Intergenic contigs are significantly over-represented in “neo” contigs (Fisher’s exact P=1.2e-20).

Additional file 2 — Table S1-S12

Table S1: Nucleotide contents of DE-kupl contigs for the TCGA LUAD dataset. Table S2: Description of event categories extracted from DE-kupl-annot tables. Table S3: General characteristics of contigs shared between LUADtcga and LUADseo. Table S4: Summary statistics for all event categories in contigs shared between LUADtcga and LUADseo. Table S5: Genes with more than 35 mapped contigs (shared LUAD contigs. Colored columns indicate ratio of contigs in said categories). Table S6: Blast results of 100 contigs mapped to IGHV genes. Table S7: Univariate Cox regression results of all categories. Table S8: Enrichment of OS-related events. Table S9: Multivariate Cox regression results of all categories. Table S10: Peptides of strong binding levels predicted by netMHCpan 4.0 from “neoRNA” contigs. Table S11: GO enrichment using host genes of immune related contigs. Table S12: Contigs correlated with the *klebsiella* contig.

Additional file 3

Correlation analysis of number of contigs and host gene expression.

