## Additional file1 Figure S1-S8 for "The contribution of uncharted RNA sequences to tumor identity in lung adenocarcinoma"

### Supplementary Figures

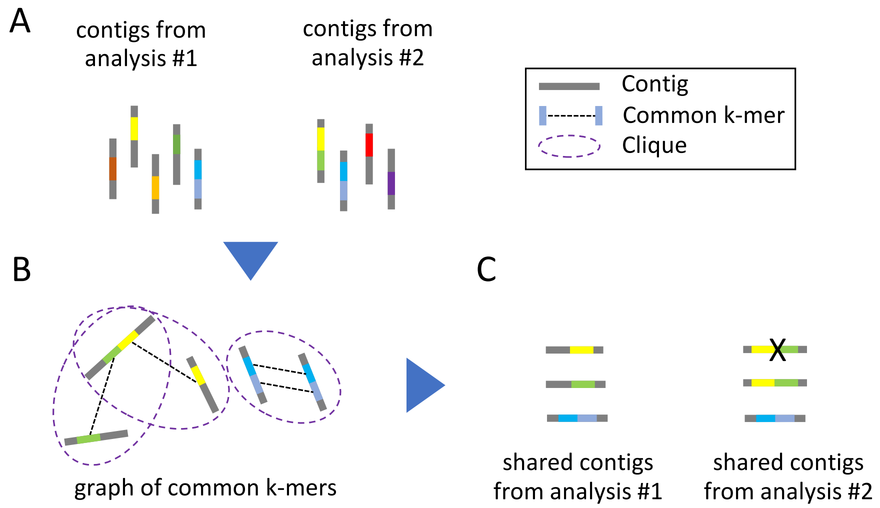


Figure S1: Protocol for detection of contigs shared between two independent datasets. (A) Contigs from each dataset. Segments of same color represent the same k-mer. (B) Cliques construction based on common k-mers. (C) Shared contigs identification based on cliques.

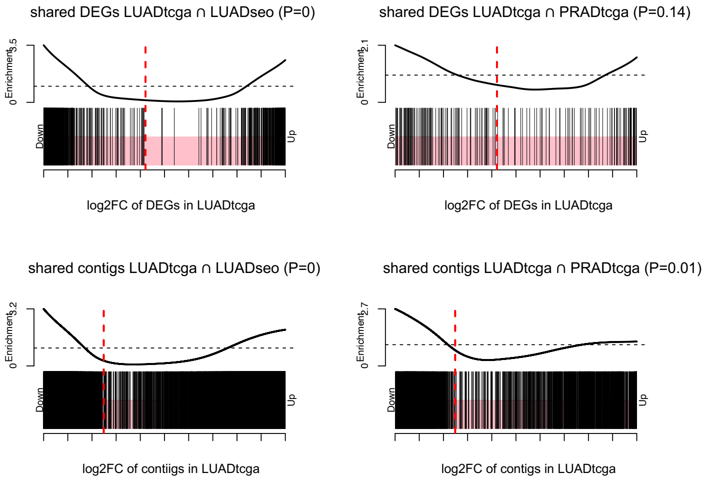


Figure S2: Enrichment analysis of shared DEGs and contigs between TCGA and SEO datasets. The x axis represents the ranked DEGs or contigs based on log2FC in ascending order. The red vertical dotted line represents the position of log2FC cutoff.

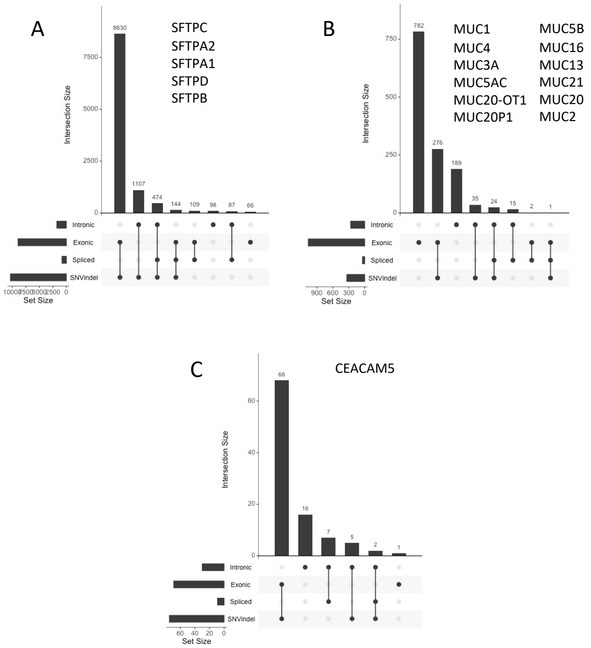


Figure S3: Contigs in hypervariable genes. Each UpsetR graph shows the intersection of contigs from categories intronic, exonic, spliced and SNV and indel.

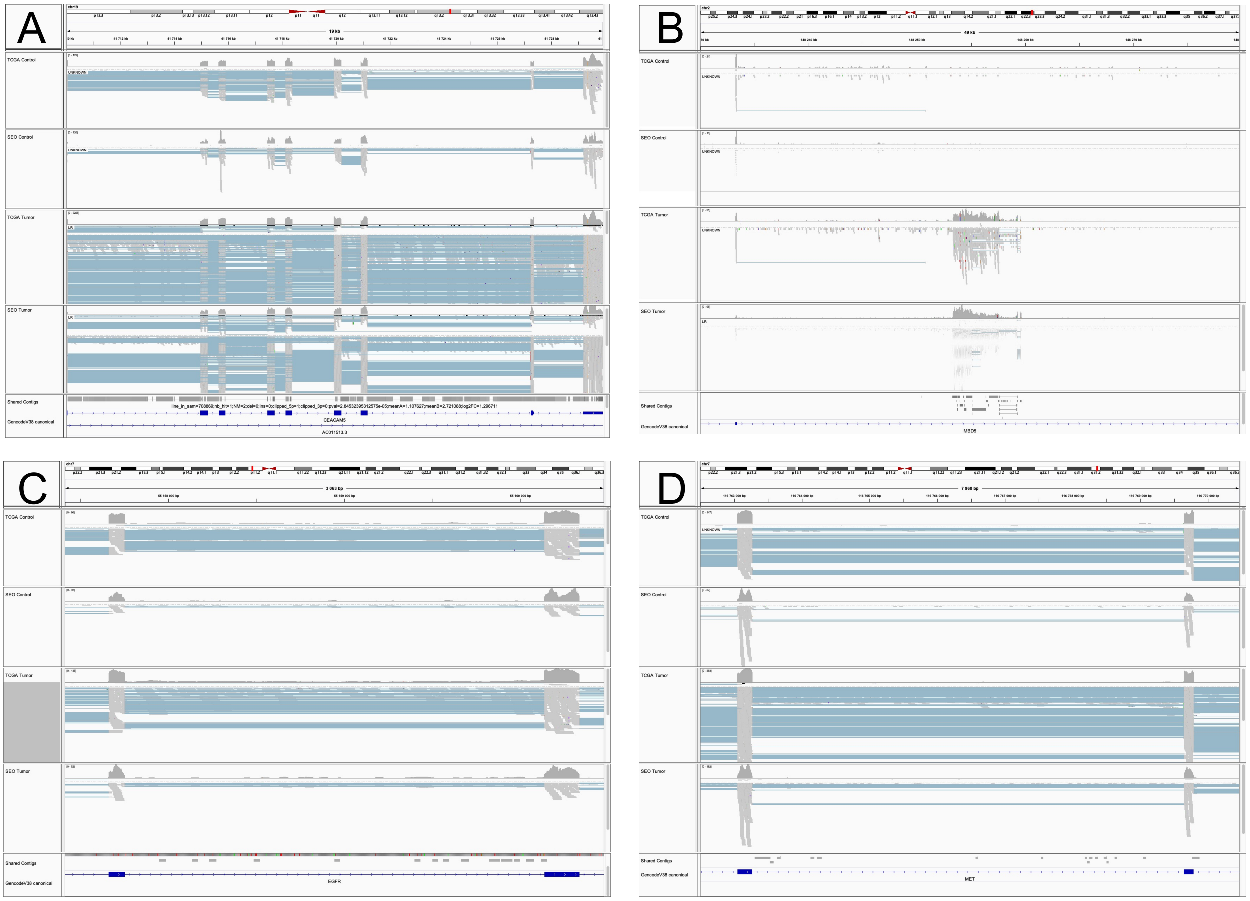

Figure S4: IGV views of intronic events. Each frame shows a metabam file composed of randomly sampled reads corresponding to the subcohort indicated on the left panel. The lower panel shows DE contigs and Gencode annotation. A: multiple intron retention in CEACAM5; B: lncRNA element expressed in an intron of MBD5; C: intron retention in EGFR; D: intron retention in MET.

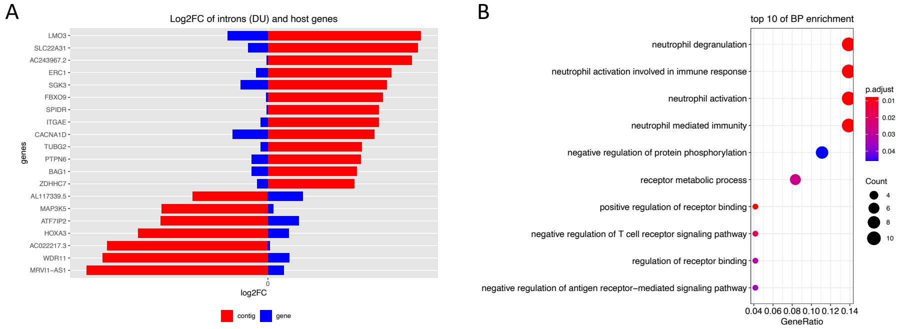


Figure S5: Intronic event analysis. (A) Log2FC values of the top 20 intronic events (DU). Red and blue colors represent the expression fold change of intronic contigs and host genes, respectively. (B) Gene Ontology functional enrichment. Color represents the P-values and size represents the ratio of genes.

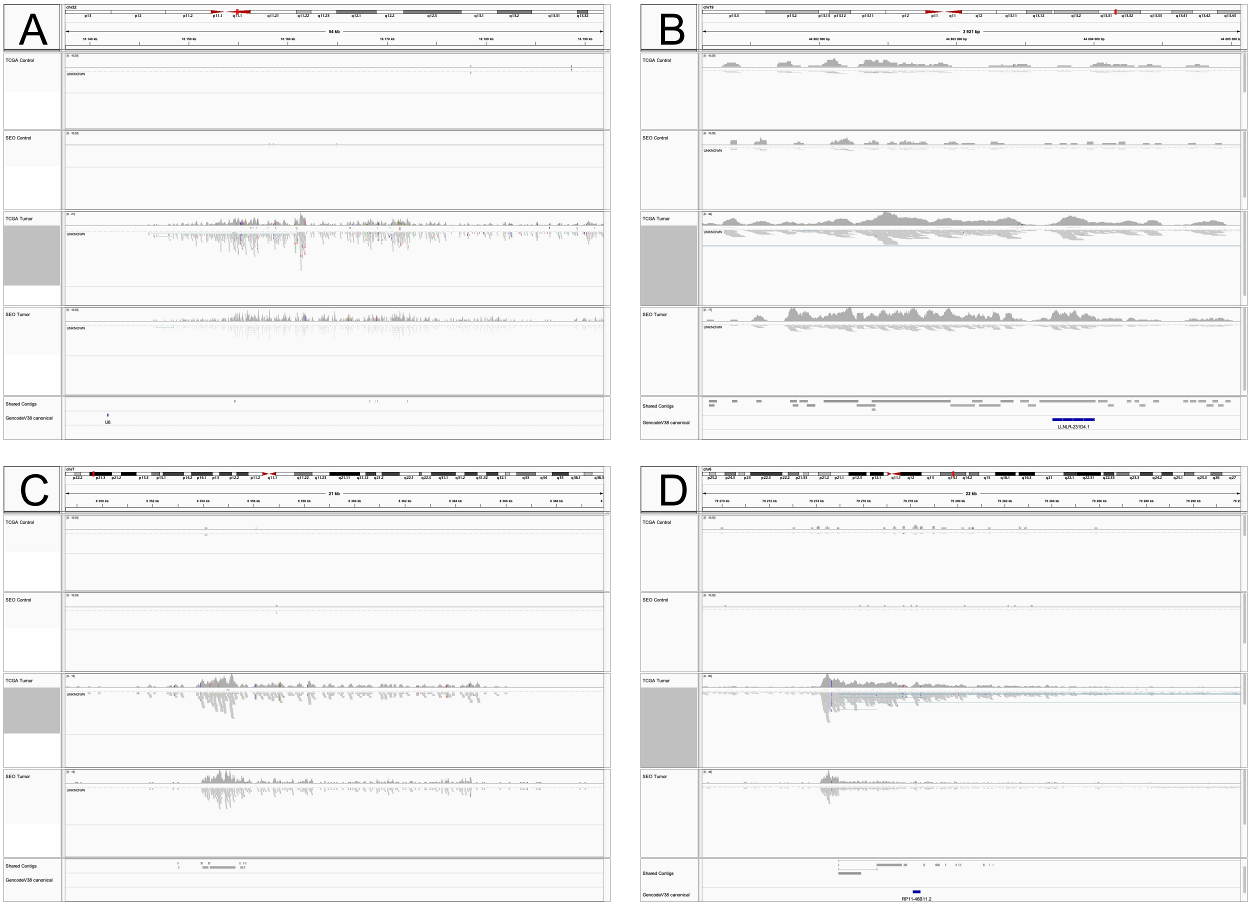


Figure S6: IGV views of lincRNA elements overexpressed in tumors. Each frame shows a metabam file composed of randomly sampled reads corresponding to the subcohort indicated on the left panel. The lower panel shows DE contigs and Gencode annotation.

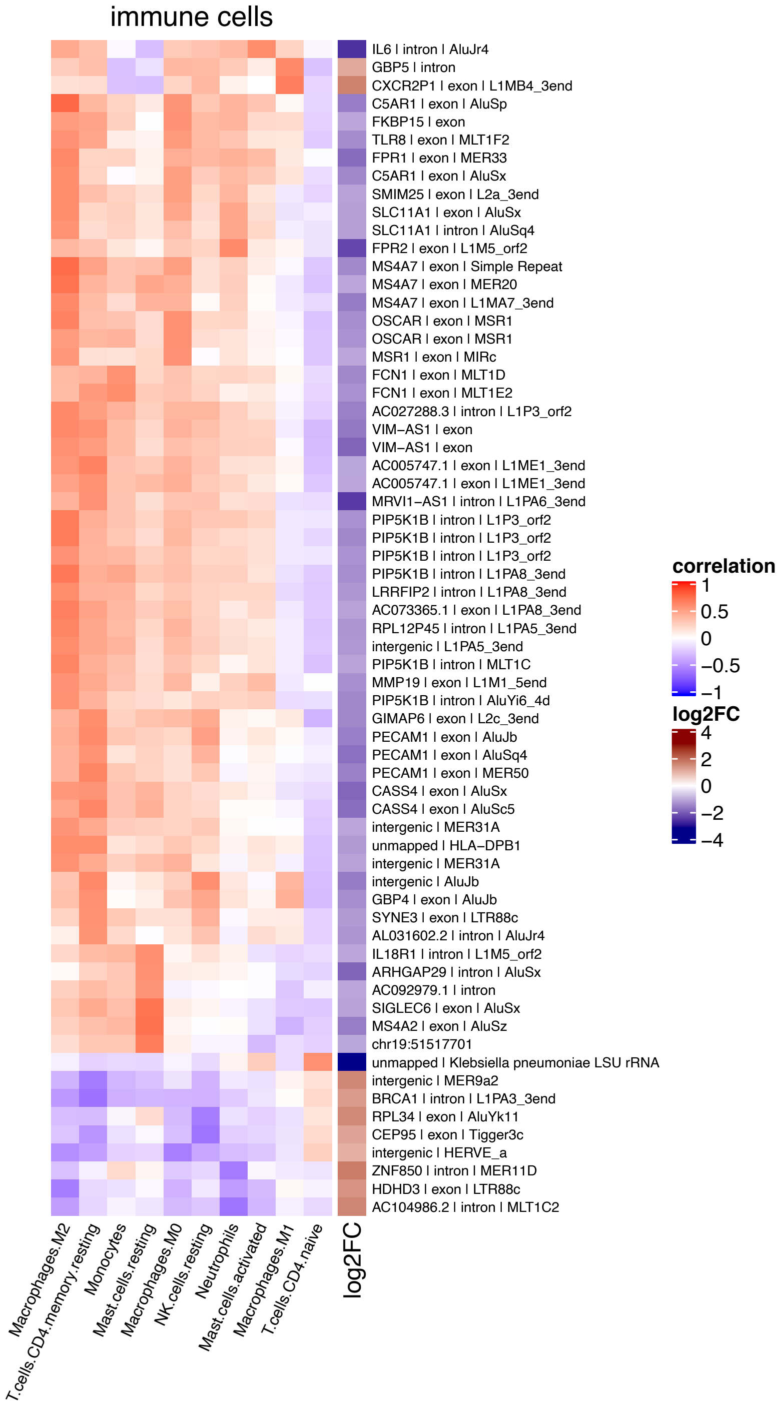


Figure S7: Heatmap of Spearman correlation coefficient (CC) of contig counts and abundance of immune cell types evaluated by CIBERSORT. All contigs with a CC>0.5 with at least one immune cell type are shown. Immune cells not correlated with at least one contig are not shown. Row names show gene symbols and repeat types of contigs, whenever applicable. Row name colors indicate different contig categories. The log2FC sidebar shows expression fold change of contigs between normal and tumor samples.

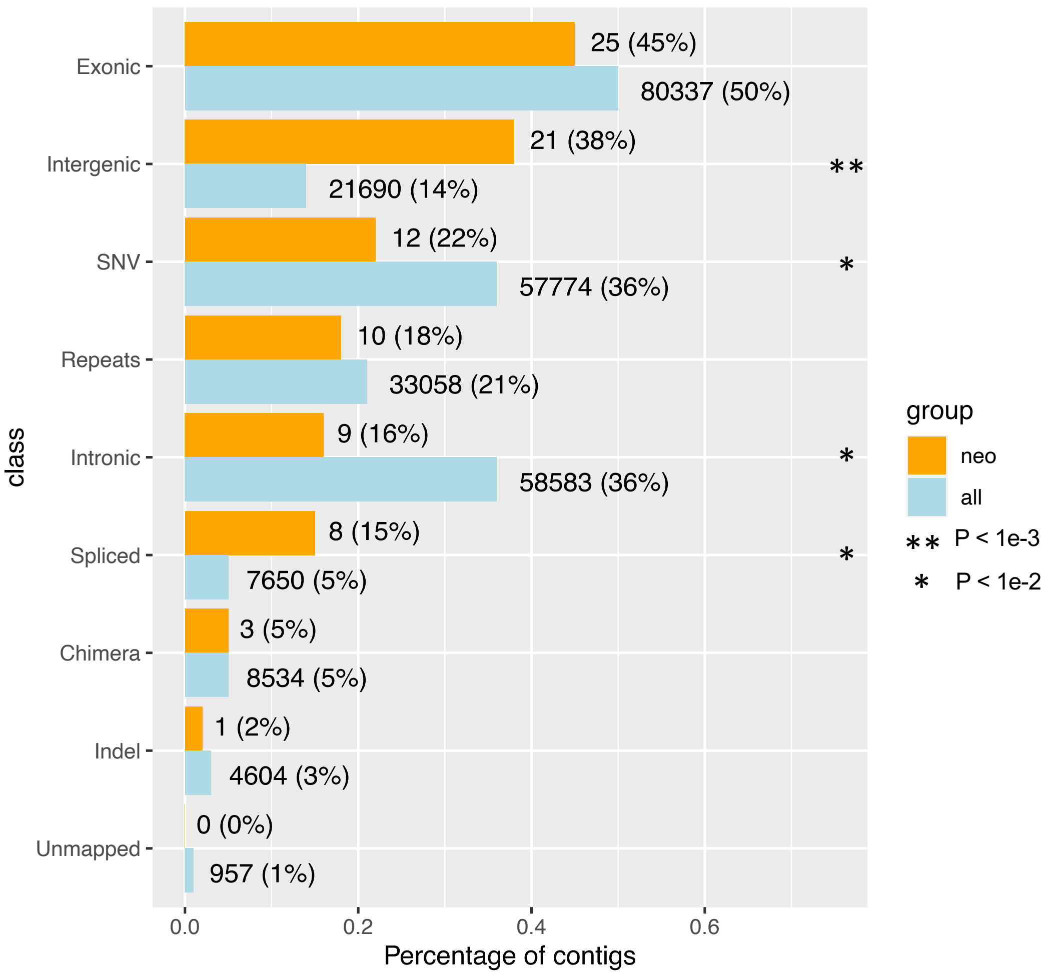


Figure S8: Types of contigs predicted to produce neoantigens ("neo", N=472) and total contigs ("all", N=2375). Intergenic contigs are significantly over-represented in "neo" contigs (Fisher's exact P=1.2e-20).
