## Additional file3 for "The contribution of uncharted RNA sequences to tumor identity in lung adenocarcinoma"

### Noise of contigs from highly expressed genes

The number of DE contigs in our analysis is at least an order of magnitude higher than the number of differentially expressed genes. One possible reason is DE-kupl identifies local events, which means each contig corresponds to one isolated event, except when two variants occur closely with a distance smaller than k. Therefore, a longer gene composed of multiple exons and introns may contribute to more differential expression contigs. Another possible reason is the high expression levels of genes. If a gene is highly expressed in tumors, it has a higher chance to introduce more variants that can be detectable by statistical methods.

To evaluate this effect, we computed correlations between numbers of contigs and expression of host genes. Fig 1A shows genes with higher expression induce more contigs. However, when considering only shared contigs, the correlation coefficient is strongly reduced (Fig 1B), suggesting shared contigs are substantially less noisy than total contigs.

On the other hand, the differential genes were selected by means of fold change. The highly expressed genes tend to have low fold change (Fig 1C). Most of these highly expressed genes can be ruled out by the fold change filtering. Therefore, the gene-level analysis is supposed to be less noisy than the contig-level analysis.

To figure out which type of event is most affected by highly expressed genes, we selected the 1% most highly expressed genes and grouped their derived contigs into different classes (Fig 1D). We found that novel repeats account for a larger proportion than the other events. The events introduced by highly expressed genes are strongly enriched in the novel repeats with a Fisher exact test P-value of 7.533e-11. The category of SNVs whose ratio is 3.1% in the pie graph is also significantly enriched (P-value = 1.459e-06). Despite a high ratio of 12.2%, the category of annotated repeats is not significantly enriched due to

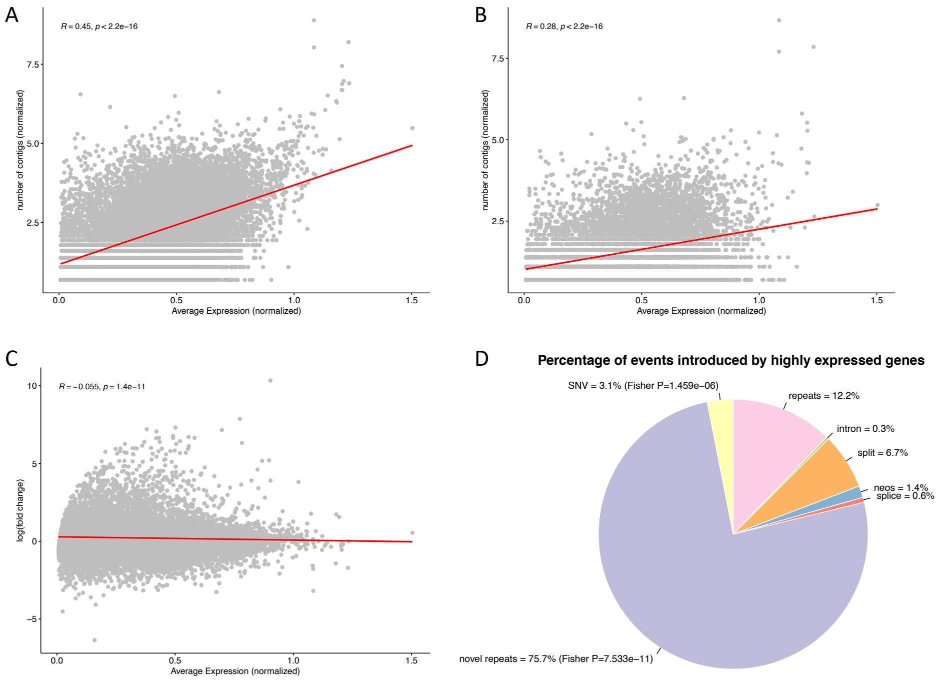


Figure 1: Noise from highly expressed genes.(A) Correlation of gene expression and numbers of contigs in the total contig list (B) Correlation of gene expression and numbers of shared contigs (C) MA-plot showing the correlation between gene expression and fold change. (D) Percentage of contigs contributed by the top 1% highly expressed genes. Gene expression are log- and size-normalized. Red lines show linear regressions. R and P-values are computed with Pearson correlation.

its large number. No matter annotated or novel repeats, they have multiple hits on the genome. Therefore it is reasonable that these repeats are correlated with highly expressed genes. From this perspective, the high expression estimates of genes may also be due to the inclusion of repeat regions, which leads to an over-estimation of gene counts.
